# Non-equilibrium antigen recognition in acute infections

**DOI:** 10.1101/2023.04.05.535743

**Authors:** Roberto Morán-Tovar, Michael Lässig

## Abstract

The immune response to an acute primary infection is a coupled process of antigen proliferation, molecular recognition by naive B-cells, and their subsequent proliferation and antibody shedding. Here we show B-cells can efficiently recognise new antigens by a tuned kinetic proofreading mechanism, where the number of proofreading steps and the characteristic rate of each step are set by the complexity of the immune repertoire. This process produces potent, specific and fast recognition of antigens, maintaining a spectrum of genetically distinct B-cell lineages as input for affinity maturation. We show that the proliferation-recognition dynamics of a primary infection can me mapped onto a generalised Luria-Delbrück process, akin to the dynamics of the classic fluctuation experiment. We derive the resulting statistics of the activated immune repertoire: antigen binding affinity, expected size, and frequency of active B-cell clones are related by power laws. Their exponents depend on the antigen and B-cell proliferation rate, the number of proofreading steps, and the lineage density of the naive repertoire. Empirical data of mouse immune repertoires are found to be consistent with activation involving at least three proofreading steps. Our model predicts key clinical characteristics of acute infections. The primary immune response to a given antigen is strongly heterogeneous across individuals; few elite responders are distinguished by early activation of high-affinity clones. Conversely, ageing of the immune system, by reducing the density of naive clones, degrades potency and speed of pathogen recognition.

## Introduction

B-cells are a central part of the human adaptive immune system. These cells recognise antigens by specific binding: B-cell receptors (BCRs) located in the cellular membrane bind to antigenic epitopes, which are cognate binding loci on the surface of pathogens. To capture a wide range of a priori unknown pathogens, humans produce a large and diverse naive B-cell repertoire, estimated to contain about *L*_0_ ∼ 10^9^ lineages with distinct BCR genotypes [1, 2, 3] and a comparable number of circulating naive cells [4]. An acute infection is characterised by rapid, initially exponential growth of an antigen population, often starting with few particles and reaching peak densities of order 10^8^ml^−1^ within a few days [5, 6, 7, 8]. At some stage of this process, free antigens start to bind to B-cells in lineages of sufficiently high binding affinity. Antigen binding can activate B-cells, inducing rapid proliferation and shedding of free antibodies that eventually clear the antigen. Activated B-cells also form germinal centres and create immunological memory [9, 10]. Primary infections are estimated to generate multiple activated B-cell lineages, *L*_act_ ≳ 10^2^ [11].

The exponential growth of antigens, together with a large number of B-cell lineages, generates a formidable, real-time specificity problem for recognition. Consider an antigen that activates a high-affinity B-cell lineage at a given point of time. One day later, at a > 100fold higher population density, the antigen can potentially activate a large number of low-affinity lineages, generating a poor overall response of the immune repertoire. The actual process activates only a tiny fraction of the pre-infection repertoire. Previous work has established upper bounds of order 10^−5^ [12, 13], and a lower bound follows from recent data, *L*_act_/*L*_0_ ≳ 10^−7^ [11]. How is such highly specific immune response possible?

To address this problem, we develop a model for the recognition dynamics that consists of three steps: antigen proliferation, molecular activation, and subsequent proliferation of activated B-cells (Fig. 1). The B-cell activation process contains kinetic proofreading: a series of multiple, thermodynamically irreversible steps [14, 15]. In an appropriate regime of rate parameters, processes with kinetic proofreading are known to increase the affinity discrimination of their output compared to near-equilibrium processes. For the activation process of Fig. 1, we show that the activation rate of weak binding B-cells depends on the antigen-BCR binding constant and on the number of activation steps, *u*_act_ ∼ *K*^−*p*^.

**Fig. 1.**
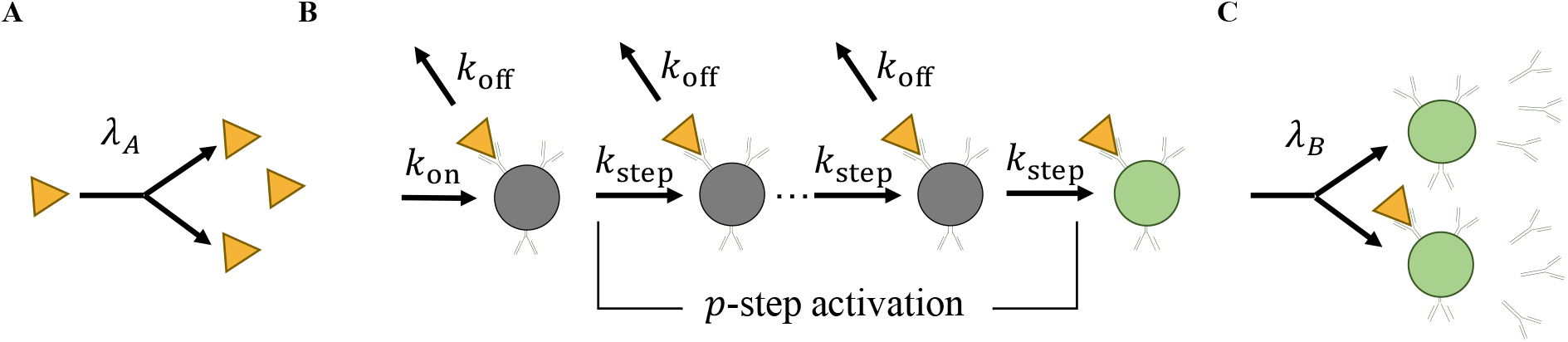
Antigen recognition in a primary infection. In a minimal model, immune recognition of a new antigen involves three stages. **(A)** Exponential replication of free antigens with rate *λ*_*A*_. **(B)** Activation of naive B-cells: binding of antigens with association rate *k*_on_, dissociation rate *k*_off_, and *p* irreversible activation steps with rate *k*_step_. This mechanism yields a total activation rate *u*_act_ that decays as *u*_act_ ∼ (*k*_step_/*k*_off_)^*p*^ in the low-affinity regime (see text). **(C)** Exponential replication of activated B-cells and proliferation of free antibodies with rate *λ*_*B*_.

Kinetic proofreading has been recognised as a key step in the activation of T-cell immunity [16, 17, 18, 19]. For B-cells, evidence for proofreading comes from experimental observations of characteristic time lags in activation, but little is known about the underlying molecular mechanism [20]. Mechanisms observed in immune cell activation by membrane-bound antigens include BCR clustering, quorum sensing, and membrane spreading-contraction [21, 22]. Such mechanisms may contribute to proofreading, but the relevance of particular mechanisms for the specificity problem addressed in this article remains unclear. Here we use a minimal *p*-step model of activation to show that proofreading is essential for specific and timely recognition of pathogens in acute primary infections.

To understand how activation and proofreading act in the face of an exponentially increasing input signal, we map the recognition dynamics onto a generalised Luria-Delbrück process. This resembles the proliferation-mutation dynamics of the classical Luria-Delbrück experiment [23]: the antigen corresponds to the wild-type, activation to mutation, and B-cell lineages to mutant cell lineages. The new feature of the infection dynamics, which has no analogue in the original Luria-Delbrück process, is that each B-cell lineage has a specific antigen binding constant *K*. This sets the density of B-cell lineages available for activation, 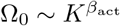, and modulates their activation rate.

In the first part of the paper, we develop the theory of generalised Luria-Delbrück processes for antigen recognition, and we compute the distribution of lineages in activated B-cell repertoires. In the second part, we turn to biological implications of the recognition dynamics. We show that proofreading is required for effective antigen recognition. Our model predicts efficient immune responses, tuned to a balance between speed and potency, at an intermediate number of proofreading steps. Recent data of mouse immune repertoires [11] are shown to be consistent with this prediction.

Activated immune repertoires of different hosts responding to the same antigen show giant fluctuations, similar to mutant populations in a classical fluctuation experiment. Such fluctuations are a hallmark of Luria-Delbrück processes [23, 24, 25, 26]. In a primary immune response, giant fluctuations are generated by “jackpot” clones of large size and high antigen affinity. We derive the underlying statistics of activated repertoires and infer clinically important characteristics of acute primary infections.

## Results

### Antigen recognition dynamics

#### Antigen-BCR binding

In the initial phase of an infection, the antigen population grows exponentially with a rate *λ*_*A*_ (Fig. 1A, 2A). For viral pathogens, this process starts with few antigen copies and reaches densities *ρ*_*A*_(*t*) of order 10^8^/ml within about 5 days, which implies replication factors > 100/d [5, 6, 7, 8]. Free antigens bind to circulating naive B-cells with rate *u*_on_(*t*) = *b*_0_*k*_on_*ρ*_*A*_(*t*) per cell, where *b*_0_ is the number of BCR per cell and *k*_on_ is the association rate (Fig. 1B). Association is known to be diffusion-limited with typical rates *k*_on_ ∼ 10^6^ M^−1^s^−1^ [27] and B-cells are known to have up to *b*_0_ = 10^5^ BCRs in their membrane [1]. Therefore, differences in antigen binding affinity between different B-cell lineages result primarily from differences in the dissociation rate *k*_off_. Human B-cells have dissociation rates in the range *k*_off_ = 10^−5^ −10^1^s^−1^ [1]. The corresponding equilibrium binding constant, *K* = *k*_off_/*k*_on_, varies in the range *K* = 10^−11^ − 10^−5^M. This constant is related to the lineage-specific energy gap between the bound and the unbound state, *K* = *K*_0_ exp(Δ*E*), where *K*_0_ is a normalisation constant and all energies are measured in units of *k*_*B*_*T* (Materials and methods). Importantly, with these parameters, the fraction of antigen-bound B-cells remains small throughout the infection process.

#### B-cell activation

Upon binding, we assume that Bcells undergo a series of *p* thermodynamically irreversible steps to activation (Fig. 1B). That is, cells in each intermediate state transform to the next state (with rate *k*_step_) or unbind from the antigen (with rate *k*_off_), but do not revert to the previous state. The stepwise, stochastic activation dynamics is an inhomogeneous Poisson process with an output activation rate *u*_act_(*t*). In the relevant regime of low antigen concentration, this rate takes the form

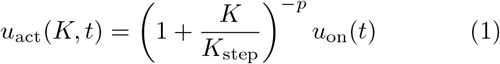

with *K*_step_ = *k*_step_/*k*_on_. In the low-affinity regime, the activation rate has the asymptotic form *u*_act_(*K, t*) ∼ (*K/K*_step_)^−*p*^ ∼ (*k*_step_/*k*_off_)^*p*^, which can be read off from Fig. 1B: each activation step generates a factor (*k*_step_/*k*_off_) relating the thermodynamic weights of consecutive intermediate states.

Next, we compute the activation probability of a B-cell lineage up to time *t, R*(*K, t*). For exponential antigen growth, we find

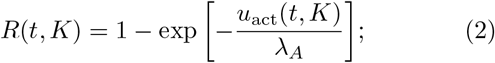

(Materials and methods). This equation describes a moving front of deterministic activation, *K*_1_(*t*) = *K*_step_ exp[(*λ*_*A*_/*p*)(*t* − *t*_1_)], where *R* reaches values of order 1 (Fig S1). The front starts at affinity 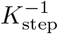 and time *t*_1_ = log[(*λ*_*A*_𝒩)/(*k*_on_*b*_0_)]/*λ*_*A*_, where 𝒩 is Avogadro’s number. With increasing antigen concentration, it moves towards lineages of decreasing antigen affinity at a *p*-dependent speed (Fig. 2B). Ahead of the front, for *K* ≫ *K*_1_(*t*), activation of individual lineages is a rare stochastic event. For *p* = 1, activation is asymptotically proportional to the inverse equilibrium constant, or Boltzmann factor, *R* ∼ *K*^−1^ ∼ exp(− Δ*E*). For *p* > 1, the non-equilibrium dynamics of kinetic proofreading generate stronger suppression of activation for weak binders, *R* ∼ *K*^−*p*^ [14, 15]. Kinetic proofreading appears to be the simplest mechanism to generate deterministic activation of high-affinity lineages together with strong suppression of low-affinity lineages; mechanisms with reversible antigen-receptor binding have *R* ∼ *K*^−1^ or remain in the stochastic regime (*R* ≪ 1) under the physiological conditions of an early primary infection (Fig. S1, Materials and methods).

**Fig. 2.**
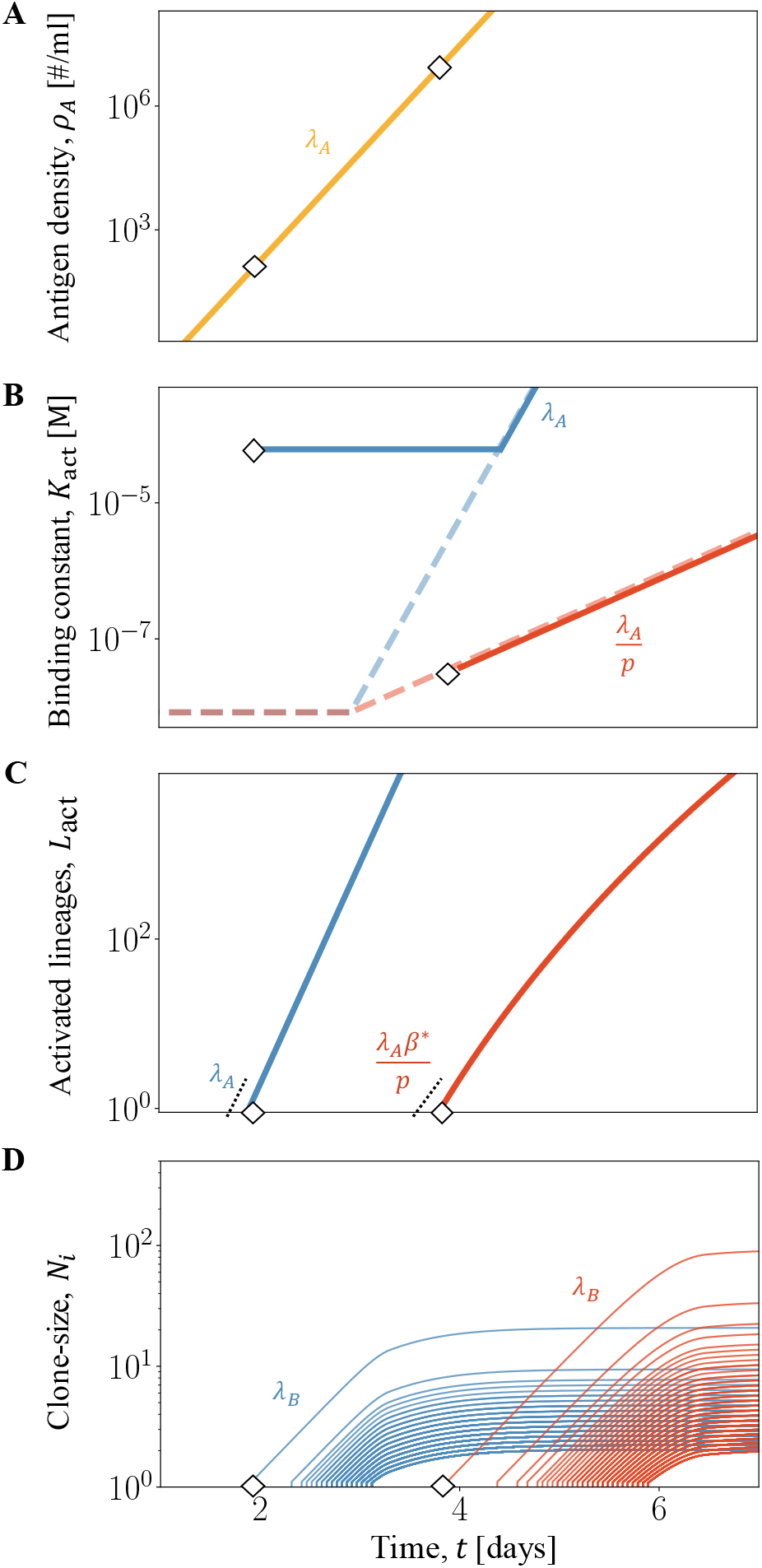
Generalised Luria-Delbrück replication-activation dynamics. **(A)** The antigen population, *N*_*A*_(*t*), grows exponentially with rate *λ*_*A*_. Diamonds mark the start of B-cell activation in the low-specificity (LS) and the high-specificity (HS) regime. **(B)** The average binding constant of activation, *K*_act_(*t*), is initially constant (LS, blue) or increases with rate *λ*_*A*_/*p* (HS, red). **(C)** The number of activated lineages, *L*_act_(*t*), grows exponentially with initial rates *λ*_*A*_ (LS) and *β*_act_*λ*_*A*_/*p* (HS). **(D)** The population of activated lineages, *N*_*i*_(*t*) (*i* = 1, 2, …,), shows initially exponential growth with rate *λ*_*B*_. In the mapping to a classical Luria-Delbrück process, the antigen population corresponds to the wild-type cell population, activation to mutation, and B-cell linages to mutant cell clones under selection. The generalised Luria-Delbrück process couples growth to neutralisation function and repertoire complexity, setting independent growth rates of *K*(*t*) and *L*(*t*). Analytical results are shown for the following parameters: activation steps, *p* = 1 (LS), *p* = 4 (HS); kinetic parameters, *k*_on_ = 10^6^*M* ^−1^*s*^−1^, *k*_step_ = 0.5min^−1^; number of BCRs per cell, *b*_0_ = 10^5^; growth rates *λ*_*A*_ = 6d^−1^, *λ*_*B*_ = 2d^−1^; repertoire size *L*_0_ = 10^8^; carrying capacity 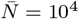.

Lineage activation marks the onset of the immune response to a new antigen. Activated cells proliferate exponentially with an initial rate *λ*_*B*_ that is comparable to *λ*_*A*_ [28] and shed free antibodies that can neutralise antigens (Fig. 1C). As the activated repertoire grows, cells start to compete for space and resources, including T-cell help [29]. Here we model the clone dynamics as logistic growth, 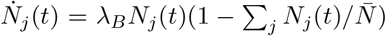, where *N*_*j*_(*t*) is the number of activated cells in clone *j* and 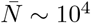 [28] is a carrying capacity for the total size of the activated repertoire (Fig. 2D; here and below, overbars refer to the repertoire statistics at carrying capacity).

#### Repertoire response to a given antigen

To characterise the immune repertoire available for primary response against a given antigen, we grade naive B-cell lineages by their affinity to the antigen’s binding locus (epitope). We use a simple sequence-specific binding energy model, where epitopes and their cognate BCR are sequence segments, **a** = (*a*_1_, …, *a*_*ℓ*_) and **b** = (*b*_1_, …, *b*_*ℓ*_). Binding aligns these segments and couples pairs of aligned amino acids, and the binding energy gap Δ*E*(*K*) = log(*K/K*_0_) is additive,

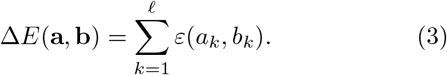

For a given antigen, this model defines the density of naive lineages available for activation, Ω_0_(*K*) = (*K*d/d*K*)*L*_0_(*K*), where *L*_0_(*K*) is the expected number of lineages in an individual with binding constant < *K* to the epitope **a** (Fig. 3). Here we assume that naive repertoires are randomly sampled from an underlying amino acid distribution [3]. Hence, most lineages bind only weakly to a new antigen (*K* ∼ 0). The expected minimum binding constant in an individual, *K*\*, is given by the condition *L*_0_(*K*\*) = 1. This point is to be distinguished from the global minimum for a given antigen, *K*_*m*_, which often corresponds to a unique BCR genotype **b**_*m*_ (called the Master sequence). Because individual repertoires cover only a small fraction of the BCR genotype space, the expected maximum antigen affinity remains substantially below the Master sequence (*K*\* > *K*_*m*_). To characterise the strong-binding tail of the lineage spectrum, we define the micro-canonical entropy, *S*(*K*) = logΩ_0_(*K*) + const., and the associated inverse reduced temperature,

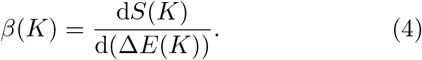

This function measures the exponential increase in lineage density with binding energy, Ω_0_(*K*′) ∼ exp[*β*(*K*) Δ*E*(*K*′)], in the vicinity of a given point *K*. As we will show below, the inverse temperature at maximum binding, *β*\* = *β*(*K*\*), is a key determinant of the activation dynamics.

**Fig. 3.**
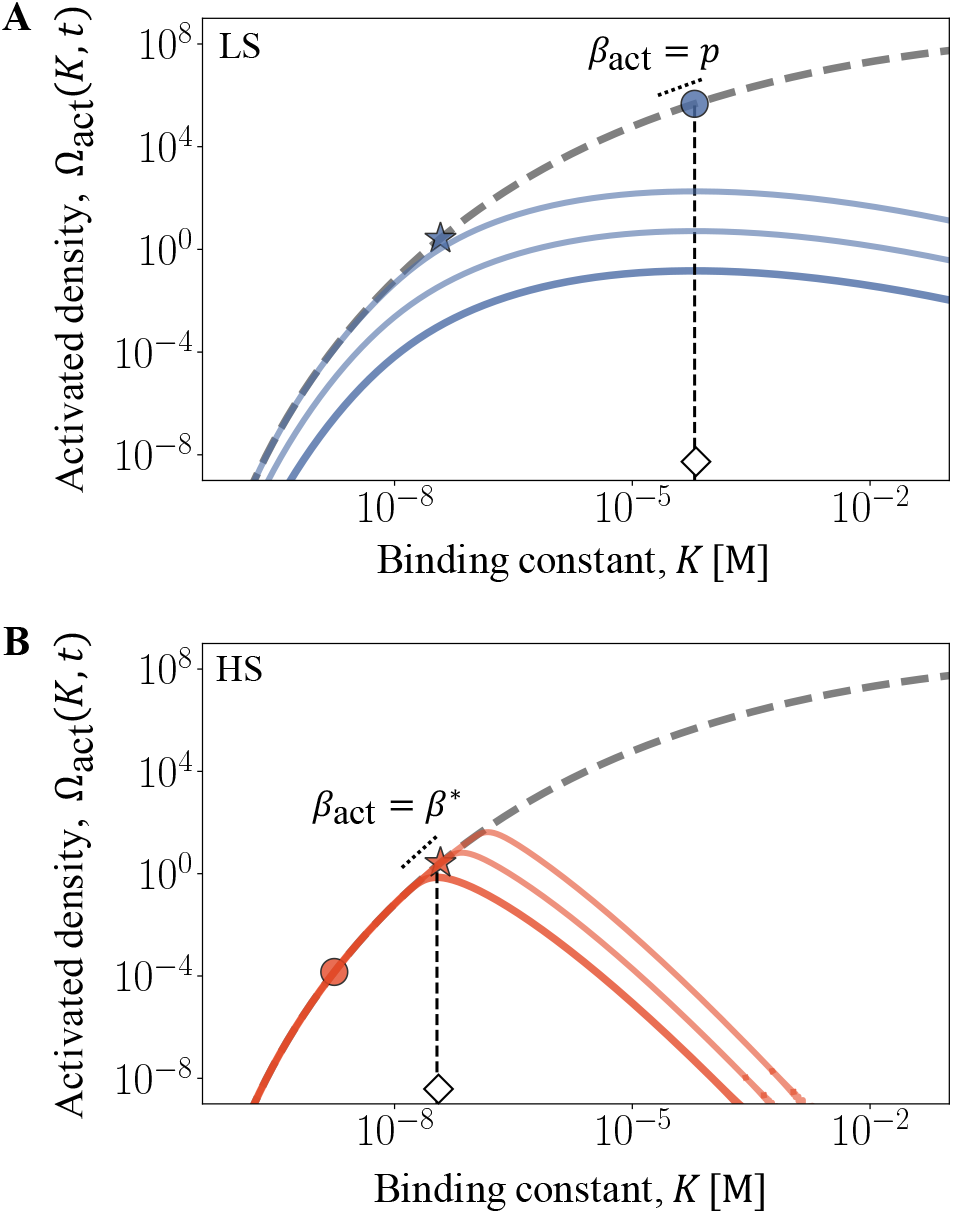
B-cell response repertoires. Density of naive B-cell lineages, Ω_0_(*K*) (gray); density of activated lineages, Ω(*K, t*), at the start of activation(*t* = *t*_act_, thick) and at two subsequent time points (thin) in the **(A)** LS regime (*p* = 1) and **(B)** HS regime (*p* = 4). Specific values of *K* are marked on the spectral function: *K*_*p*_ (dot); strongest antigen binding in typical individuals, *K*\* (star); start of activation, *K*_act_ (open diamond) with initial inverse activation temperature, *β*_act_ (dotted line). Energy model: TCRen; other parameters as in Fig. 2.

#### Repertoire response to diverse antigens

How comparable are the response repertoires of different antigens? To address this question, we evaluate the lineage spectrum Ω_0_(*K*) for a random sample of antigenic epitopes **a** (Fig. S2). The interaction energy matrix *ε*(*a, b*) of our main analysis is proportional to the TCRec matrix *ε*(*a, b*) originally inferred for T cell receptors [30]; similar spectra are obtained from the Miyazawa-Jernigan matrix [31] and from normally distributed random energies. For a given antigen, the spectral density depends on broad statistical features of the energy matrix, including the binding length *ℓ* and the variance of interaction energies, 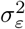 (Materials and methods). Here we determine these parameters from observed binding constants *K*\* ∼ 10^−7^M and *K*_*m*_ ∼ 10^−11^M of high-affinity antibodies generated in primary infections and of ultra-potent antibodies, respectively [32, 1].

Remarkably, these physiological constraints generate a consistent ensemble of response repertoires. First, the distributions of inferred binding lengths and of the rms. energy variation per site are strongly peaked around values *ℓ* ∼ 20 and *σ*_*ε*_/(*ℓ*^1/2^) ∼ 1, respectively, which are in tune with known examples (Fig. S2). Second, the lineage densities Ω_0_(*K*) depend only weakly on the antigen sequence **a** and have a nearly universal shape (Fig. S2). In other words, the antigen-averaged spectral density Ω_0_(*K*) captures the response repertoire available in a typical primary infection. In particular, response repertoires of different antigens with similar *K*\* and *K*_*m*_ have similar inverse activation temperatures, *β*\* = (2.5 *±* 0.3).

#### Repertoire activation

The spectral density of naive lineages and the recognition function *R*(*K, t*) determine the time-dependent density of activated lineages,

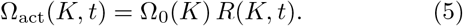

In Fig. 3, we plot Ω_act_(*K, t*) at subsequent times for two different numbers of activation steps, with and without proofreading (*p* = 1, 4). 4AB shows simulations of the stochastic B-cell clone population dynamics in a typical individual host. The spectral density of activated lineages is strongly peaked; its high-affinity flank is given by the density of naive lineages, its low-affinity flank by the proofreading-dependent activation dynamics. This function determines two repertoire summary statistics: the expected number of activated lineages, *L*_act_(*t*) = ∫ Ω_act_(*K, t*) d*K/K*, and their average binding constant *K*_act_(*t*) = ∫ *K*Ω_act_(*K, t*) d*K/K*. Activation starts at an expected time *t*_act_ given by the condition *L*_act_(*t*_act_) = 1. This sets the initial binding constant *K*_act_ ≡ *K*_act_(*t*_act_) (marked by diamonds in Fig. 3) and the inverse activation temperature *β*_act_ ≡ *β*_act_(*K*_act_) (marked by a tangent line). Importantly, activation has two dynamical regimes.

In the *low-specificity* (LS) regime, for small values of *p*, activation is peaked on the low-affinity flank of the spectral function (Fig. 3A). The starting point *K*_act_ = *K*_*p*_ is determined by the condition *β*_act_ = *p*, which follows from the asymptotic form *R* ∼ *K*^−*p*^ given by equation (2). In this regime, the number of activation steps, *p*, determines the specificity of recognition; the activation probability and clone size of individual lineages remains small (Fig. 4A). The LS activation dynamics is characterised by

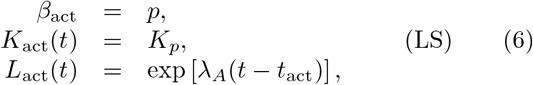

as shown in Fig. 2BC; details are given in Materials and methods. This regime ends at a crossover point *p* = *β*\*, where *K*_*p*_ reaches the expected minimum binding constant, *K*\*.

**Fig. 4.**
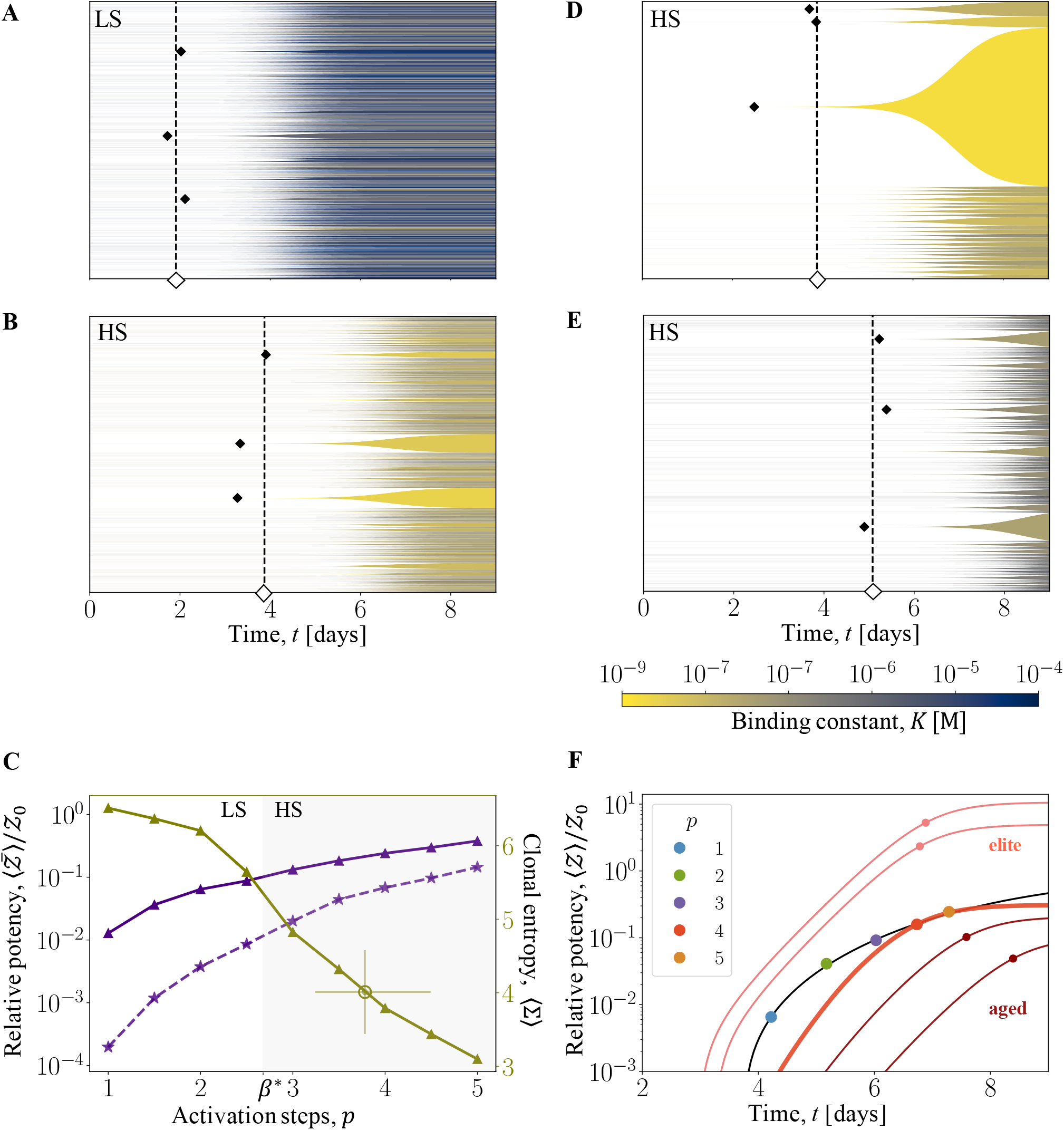
Proofreading determines potency, speed, and diversity of recognition. **(A, B)** Activated B-cell repertoires in randomly sampled individuals in the LS regime (*p* = 1) and in the HS regime (*p* = 4). Muller plots with filled areas representing individual clones (height: time-dependent population size, shading: antigen binding constant, *K*, as given by color bar); activation times of the first few clones (filled diamonds) are shown together with the expectation value *t*_act_ (open diamond). **(C)** Antiserum potency, 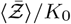, potency component of the largest clone, 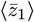, and Shannon entropy of the activated repertoire, σ. Population averages are shown as functions of the number of activation steps, *p*, at carrying capacity and relative to a reference value *Ƶ*_0_ = 1M. Shading marks the HS regime (*p* ≳ *β*\*), an open circle with error bars gives the empirical entropy inferred from the repertoire data of ref. [11]. **(D)** Activated B-cell repertoire in an elite neutraliser occurring at population frequency 10^−3^. This repertoire is marked by the early activation of a high-affinity “jackpot” clone. **(E)** Activated B-cell repertoire of an aged individual with a 10x reduced number of naive lineages. The onset of activation, *t*_act_ (black dashed line), is delayed with respect to a full repertoire (panel B). **(F)** Time-dependent antiserum potency, ⟨*Ƶ*⟩(*t*). The population average is shown in the HS regime (*p* = 4, thick red line); the half-saturation point (*t*_50_, *Z*_50_) is marked by a dot. The family of half-saturation points for different values of *p* characterises the tradeoff between potency and speed of typical immune responses, which defines a Pareto line (black line). The potency *Ƶ*(*t*) of typical elite neutralisers (at population frequencies 10^−3^ and 10^−4^, light red lines) is above, that of aged immune systems (at 10x and 100x lineage reduction, dark red lines) is below the population average (HS regime, *p* = 3). Model parameters as in Fig. 2.

In the *high-specificity* (HS) regime, for *p* > *β*\*, activation starts at *K*_act_ = *K*\* and *β*_act_ = *β*\*, then follows the deterministic front *K*_act_(*t*) = *K*_1_(*t*) (Fig. 3B). Hence, lineages are activated deterministically and in order of decreasing antigen affinity. In this regime, the spectral density of the naive B-cell repertoire determines the specificity of recognition; high-affinity lineages reach substantial clone size (Fig. 4B). We find in the HS activation dynamics

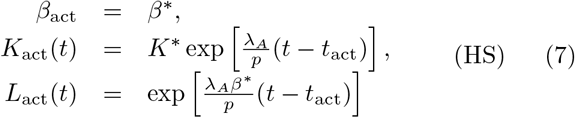

as shown Fig. 2BC; details are given in Materials and methods. In the next section, we will show that these regimes generate drastically different immune responses. We note that the growth rates *λ*_*A*_/*p* and *β*\**λ*_*A*_/*p* in equation (7) are specific to generalised Luria-Delbrück processes as defined in this paper (Fig. 2). The analogue of *L*_act_(*t*) in a classical Luria-Delbrück process is the number of mutant clones. Given a constant molecular clock of mutations, this number always grows with rate *λ*_*A*_, proportionally to the wild-type population size.

### Biological consequences

#### Efficient immune response by kinetic proofreading

As shown in Fig. 3, kinetic proofreading in the HS regime generates a specific profile of B-cell lineages generated in a primary infection: deterministic activation of high-affinity lineages is coupled to strong suppression of low-affinity lineages. To quantify the impact of this profile on immune function, we evaluate the antiserum potency

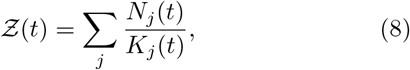

as well as the contributions of individual lineages, *z*_*j*_(*t*) = *N*_*j*_(*t*)/*K*_*j*_ (the index *j* = 1, 2, … orders clones by decreasing size). The function Ƶ(*t*) sums the antigen affinities of all activated cells and determines the fraction of neutralised virions; it reaches a saturation value 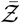 at carrying capacity of the activated repertoire. We measure potency relative to a reference value 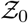 describing a hypothetical repertoire with homogeneous binding constant *K*\*. In Fig. 4C, we plot the population average 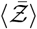 as a function of the number of proofreading steps (here and below, population averages are denoted by brackets). Potency comes close to the reference value in the HS regime, but quickly drops with decreasing *p* in the LS regime. Without proofreading (*p* = 1), 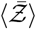 is about 20fold lower than at *p* = 4, argued below to be the approximate number of proofreading steps in human B-cell activation. The difference between activation regimes is even more pronounced for the potency contribution of the largest clone, 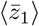 (Fig. 4C). In the HS regime, where the largest clone is likely also the clone of highest affinity, 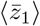 contributes a substantial fraction of the total potency. There is again a rapid drop in the LS regime; without proofreading, 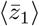 is about 1000fold lower than at *p* = 4.

Another striking difference between the activation regime is in the activation probability of individual lineages, as given by the recognition function *R*(*K, t*). In the HS regime, almost all high-affinity lineages get activated (*R* reaches values close to 1); in the LS regime, most available highaffinity lineages do not get activated and are waisted for pathogen suppression (*R* ≪ 1). Together, we conclude that the lineage activation profile generated by kinetic proofreading in the HS regime is a prerequisite for a potent, specific, and efficient primary immune response.

#### Repertoire-tuned proofreading

Fig. 4AB shows two further functional differences between the activation regimes. In the LS regime, a large number of lineages gets activated, but these clones reach only small population frequencies at carrying capacity, 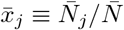. In the HS regime, activation gets increasingly focused on few high-frequency and high-affinity lineages. We can describe the diversity of activated repertoires by the Shannon entropy 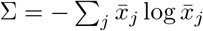. In the LS regime, the population average ⟨Σ⟩ is large and varies only weakly; in the HS regime, ⟨Σ⟩ drops substantially with increasing *p* (Fig. 4C). Subsequent to activation, a part of the B-cells undergoes affinity maturation in germinal centers. This mutation-selection process produces high-affinity plasma B-cells, as well as a diverse set of memory B-cells. In the HS regime, the larger repertoire diversity found close to the crossover point (*p* = *β*\*) serves both channels of affinity maturation: it facilitates the search for mutational paths towards high-affinity BCR genotypes in plasma cells, and it provides diverse input for memory cell formation [33].

In Fig. 4F we plot the time-dependent, population-averaged potency for different values of *p*. The response time *t*_50_, where ⟨*Ƶ*⟩(*t*) reaches the half-saturation point 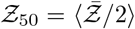, is marked by dots. The *p*-dependent increase in potency is coupled to an increased time delay of activation, caused by the sequence of intermediate steps and by the constraint to high-affinity lineages. In the HS regime, increasing *p* yields a diminishing return of potency, while *t*_50_ continues to increase proportionally to *p*. Similarly, efficient proofreading requires a sufficiently small activation rate. In the HS regime, for *k*_step_ < *K*\**k*_on_, decreasing *k*_step_ yields a diminishing return of potency, while *t*_50_ continues to increase proportionally to 1/*k*_step_ (Fig. S3). The tradeoff between potency and speed of immune response defines a Pareto surface. This tradeoff, together with the entropy pattern, suggests that optimal immune response is tuned to the spectral density of the naive repertoire: the number and rate of proofreading steps are in the HS regime close to the crossover point *p* = *β*\* and *k*_step_ = *K*\**k*_on_, respectively.

While there is no direct evidence of proofreading in B-cell activation to date, available data suggests that immune response is in the tuned regime. Specifically, using recent data of clonal B-cell populations in early germinal centres of mice [11], we infer a repertoire entropy Σ = 4.0 *±* 0.6 (circle in Fig. 4C see Materials and methods), which is consistent with proofreading in the range *p* = 3.7[3.2, 4.5]. This argument is further supported by the inferred clone size statistics, to which we now turn.

#### Clone size and affinity statistics

B-cell immune repertoires are known to have broad variation of clone sizes, which can be described by power-law distributions [34, 35, 36]. Luria-Delbrück processes provide a simple explanation for such power laws: they relate observables that depend exponentially on time (Fig. 2). First, consider the relation between clone size and probability of occurrence in an individual’s repertoire. More lineages are activated later (*L* ∼ exp[(*λ*_*A*_*β*_act_/*p*) *t*]), but these clones reach smaller size 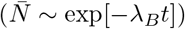. This relates clone size to rank,

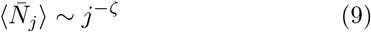

with

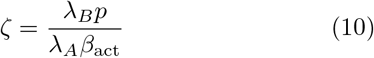

and *β*_act_ = min(*p, β*\*); see Materials and methods for details. The index *j* = 1, 2, … again orders the clones in an individual’s repertoire by size. The cumulative distribution aggregated over individuals has the form Φ(*N*) ∼ *N* ^−1/*ζ*^, which is equivalent to equation (9), and spans 3 orders of magnitude in size (Fig. S4, Materials and methods). Simulations confirm these power laws; the clone-rank statistics in a set of randomly chosen individuals follows the same pattern (Fig. 5A, S4). In the HS regime, the exponent *ζ* increases monotonically as a function of *p*, reflecting the increasing bias to large clone size generated by proofreading (Fig. 5D).

**Fig. 5.**
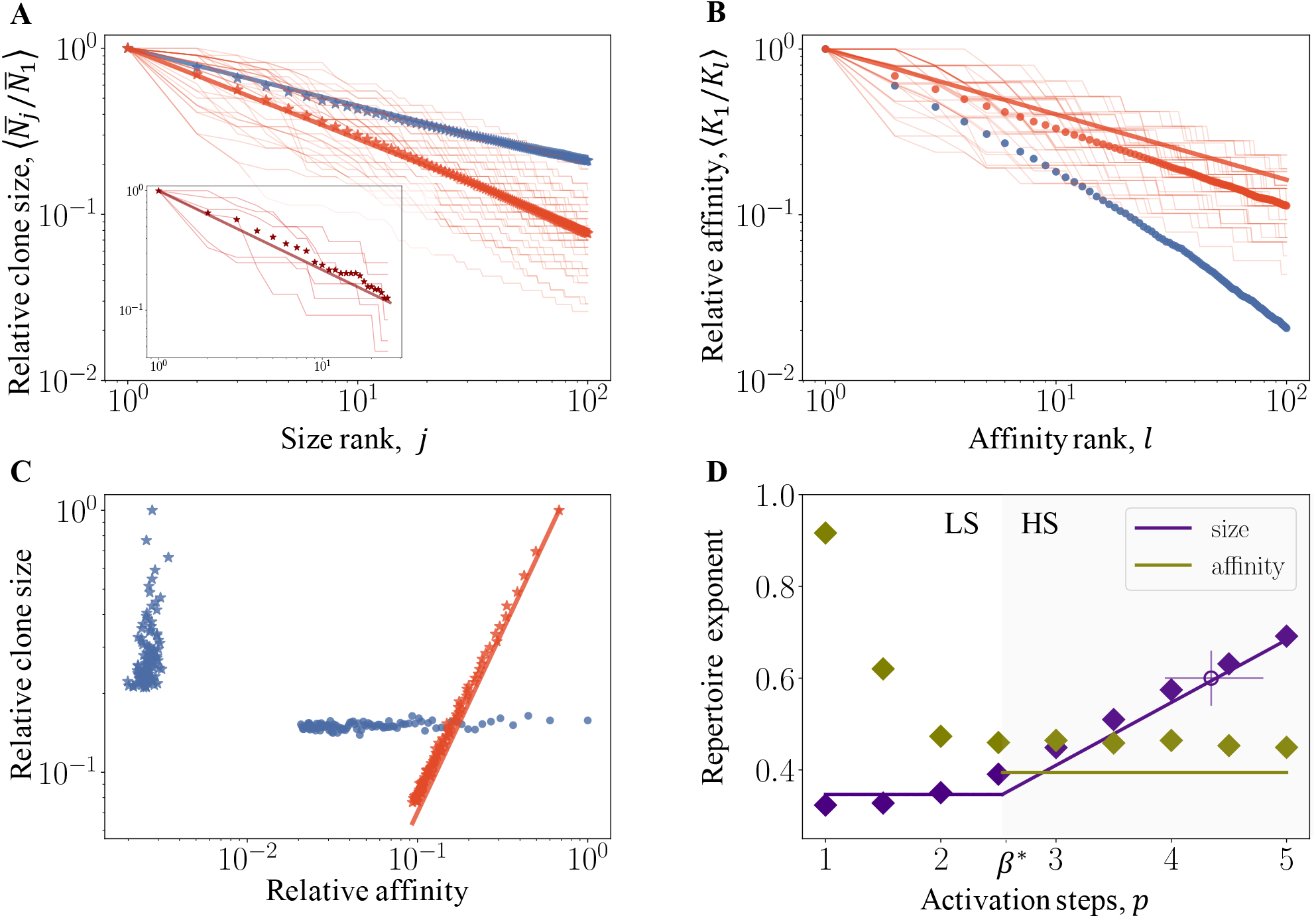
Clone size and affinity statistics of activated repertoires. **(A)** Clone size statistics. Simulation results for the average relative clone size, 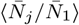, are plotted against the size rank, *j* = 1, …100, for *p* = 1, 4 (blue and red, respectively). Lines mark the expected power law, 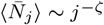, as given by equation (9). Thin lines give rankings in a set of randomly chosen individuals. Insert: empirical rank-size relation inferred from the repertoire data of ref. [11]. **(B)** Affinity statistics. Simulation results for the average relative antigen binding constant, ⟨*K*_*l*_/*K*_1_⟩, are plotted against the affinity rank, *l* = 1, …, 100. A line marks the expected power law in the HS regime, 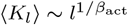, as given by equation (11). **(C)** Size-affinity correlation. HS regime: average relative affinity, ⟨*K*_*j*_/*K*_1_⟩ vs. average relative clone size, 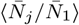 for the largest clones (*j* = 1, …, 100; red stars), together with the expected power law correlation, 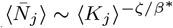 (red line). LS regime: ⟨*K*_*j*_/*K*_1_⟩ vs. 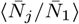for the largest clones (*j* = 1, …, 100; blue stars), and ⟨*K*_*l*_/*K*_1_⟩ vs. 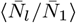for high-affinity clones (*l* = 1, …, 100; blue circles), indicating loss of the size-affinity correlation. **(D)** Empirical repertoire exponents from size and affinity rankings are plotted as functions of the number of activation steps, *p* (diamonds); the high-specificity regime is marked by shading. Lines mark power laws emerging from the Luria-Delbrück proliferation-activation dynamics; their exponents *ζ* and *β*\* depend on the proliferation rates of antigen and activated B-cells, the activation temperature, and the number of proofreading steps. An open circle with error bars gives *ζ* inferred from the repertoire data of ref. [11]. Model parameters as in Fig. 2.

To test the prediction of equation (9), we use again the mouse repertoire data reported in ref. [11]. The size-rank statistics of clones can be fit to a power law with exponent *ζ* = 0.60 *±* 0.06 (insert in Fig. 5A). The resulting estimate *p* = 4.3 *±* 0.4 (Fig. 5D) is consistent with the value inferred from the clonal entropy, confirming our inference of proofreading at an intermediate number of steps. Remarkably, clone size distributions extracted from data of human memory B-cell repertoires [37] show power laws with a similar exponent, *ζ* = 0.57 *±* 0.12 [36]. Given that memory cells are in the same affinity range than activated naive cells [38], this may point to a common dynamical origin; however, our present model contains only the primary activation dynamics and is not directly applicable to memory cells. In refs. [35, 36], the power laws of memory repertoires have been attributed to long-term selection by recurrent infections.

Generalised Luria-Delbrück processes include phenotype (here, antigen affinity) statistics and predict new power laws observable in repertoire data. In the HS regime, activation occurs on a moving front, as given by equation (7). This relates affinity to rank,

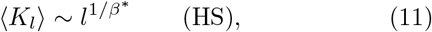

where the index *l* orders clones by decreasing affinity (Fig. 5B, Materials and methods). Equation (11) is again equivalent to a power law in the spectral density, 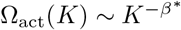, and is consistent with the affinity-rank statistics in randomly sampled individuals. The exponent 1/*β*\* equals the activation temperature of the naive repertoire, as given by equation (4). By combining equations (9) and (11), we obtain a power-law relation between size and affinity,

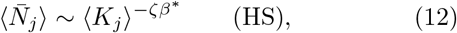

as shown in Fig. 5C. In the HS regime, the size and affinity rankings coincide up to fluctuations, because both are related to time: high-affinity clones get activated before low-affinity clones. The resulting potency-rank relation, 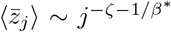, is shown in Fig. S4. In the LS regime, the size-affinity correlation is lost. Large clones have affinities of order *K*_*p*_, high-affinity clones have small size and show a faster decline of affinity with rank than in the HS regime (Fig. 3A, 5BC). Hence, empirical observations of this correlation can provide specific evidence for activation by proofreading in the HS regime.

#### Elite neutralisers

Generalised Luria-Delbrück immune activation shows particularly pronounced variation between hosts. In the HS regime, a subset of *elite neutralisers* is distinguished by early activation of a single high-affinity clone.

This jackpot clone dominates the activated immune repertoire and generates exceptionally high potency. Fig. 4D shows an example of the activation dynamics that occurs in one of 10^3^ individuals, which is to be compared with the pattern in typical individuals (Fig. 4B). Such elite neutralisers are ahead of the Pareto surface of typical immune responses (Fig. 4F). The cumulative distribution Φ(*Ƶ*), which gives the fraction of responders with saturation potency > *Ƶ*, displays two regimes of elite neutralisers (Fig. S4). In the pre-asymptotic regime, the jackpot clone takes a large part but not all of the repertoire 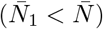. The pre-asymptotic potency distribution turns out to be dominated by clone size fluctuations, which implies Φ(*Ƶ*) ∼ *Ƶ*^−1/*ζ*^ (Materials and methods). In the asymptotic regime, the jackpot clone dominates the repertoire 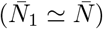; hence, Φ(*Ƶ*) is proportional to the naive density of high-affinity clones, 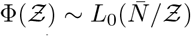. For example, one in 10^5^ individuals has a primary response with potency 100x above average, comparable to a memory immune response carried by affinity maturated B-cell lineages.

#### Immune ageing

Recent results indicate that the most prominent effect of immune ageing is a decrease in the overall size and diversity of the repertoire [39, 40, 41, 42]. Our model predicts two effects of this decrease: primary immune responses come later and with reduced potency. Simulations of the activation dynamics in an aged repertoire in the HS regime show that the activation is delayed and antigen affinities are reduced compared to a full-size repertoire (Fig. 4E, to be compared with Fig. 4B). The time-dependent potency remains behind the Pareto line and reaches a reduced value at carrying capacity (Fig. 4F). For a 10fold decrease in repertoire size, *t*_50_ increases by about 1d and 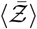 drops to half of its full-size value. Fig. S3 shows the full dependence of potency on repertoire size. Given a reference point in the HS regime, a reduction of size always has a sizeable effect, while an increase eventually induces a cross-over to the LS regime and a diminishing return of potency.

## Discussion

A potent adaptive immune response to a new antigen requires the specific activation of immune cells with high affinity to the antigen. Here we have developed a minimal, biophysical model for immune recognition by naive B-cells in a primary infection (Fig. 1). We have shown that active processes of antigen-receptor binding – kinetic proofreading – are an essential part of the initial recognition dynamics: a highly specific response to a new antigen requires proofreading with depth above a threshold value, *p* > *β*\*. Available data of mouse B-cell repertoires are consistent with proofreading close to the specificity threshold, *p* ≈ *β*\*, which amounts to at least 3 consecutive proofreading steps (Fig. 4 and 5). As shown by our model, immune responses tuned to this point are close to a functional optimum: they balance potency and speed, and they generate a diverse set of activated clones for subsequent affinity maturation.

Our model maps the proliferation-activation dynamics of a primary infection onto a generalised Luria-Delbrück process (Fig. 2). In this process, fluctuations are coupled to immune function: jackpot clones have large size and high antigen affinity, resulting in high neutralisation potency (Fig. 4). The underlying statistics of activated immune repertoires is characterised by statistical power laws with two basic exponents: the clone size exponent *ζ* and the inverse activation temperature *β*\* (Fig. 5). These exponents characterise the proofreading-dependent specificity of activated immune repertoires, as given by equations (9) and (11). Importantly, our model predicts *ζ* and *β*\* in terms of independently measurable quantities and without fit parameters. The statistics of activated B-cell repertoires shape clinically important characteristics of primary infections, including the potency drop of aged responders and the increased response of elite neutralisers (Fig. 4).

Power-law statistics of lineages and the occurrence of elite neutralisers are commonly ascribed to antigen-mediated selection on immune repertoires, often through multiple exposures [43, 35, 44, 45, 36]. Here we have shown that similar features can already emerge in the primary immune response to an acute infection, prior to any antigen-mediated selection effects. Following an acute infection, a part of the activated B-cell repertoire is further processed by affinity maturation. How this step reshapes the repertoire statistics will be studied in a subsequent publication.

Repertoire sequencing combined with neutralisation assays can test our model and probe adaptive immune systems in new ways. By recording the power-law rank-size relation of linages in early post-infection B-cell repertoires and measuring the antigen binding constant of these clones, we can extract the corresponding power laws and infer the central parameters of our model: the specificity threshold, *β*\* and the number of proofreading steps, *p* (Fig. 5). The parameter *β*\* measures the density of B-cell lineages close to the maximum-affinity lineage. This density depends on the size of the naive B-cell repertoire and on the complexity of the antigen-receptor binding motif. Both quantities emerge as key determinants of primary immune responses. In contrast, the parameter *p* characterises the molecular dynamics of antigen recognition. As we have shown, *p*-step proofreading generates an effective inverse temperature *β* ≈ *p* that measures the specificity gain in a dense repertoire. At the point of optimal recognition, *p* ∼ *β*\*, antigen recognition dynamics matches repertoire complexity. From this point, increasing *p* at a constant repertoire size *L* produces a diminishing return of proofreading; conversely, increasing *L* at constant *p* produces a diminishing return of repertoire size.

The link between repertoire size and antigen recognition machinery has implications for the macro-evolution of adaptive immune systems. The size of the total B-cell repertoire varies drastically across vertebrates, ranging from ∼ 3 × 10^5^ cells in zebrafish [34] to ∼ 10^11^ in humans [1, 2, 3]. Because potency and timeliness of immune responses are likely to be under strong selection, our model says that the functional balance of tuned proofreading, *p* ≈ *β*\*, is also a maximum of fitness. Evolutionary changes of the repertoire size are then predicted to occur together with changes of the recognition machinery, which includes number and rate of proofreading steps. Tracing these co-evolutionary dynamics by comparative cross-species studies may provide an avenue to better understand the evolution of complex immune systems.

## Acknowledgements

We thank D. Valenzano and A. Ryabova for discussions, M. Karmakar and M. Meijers for a careful reading of the manuscript, and all members of the Lässig lab for input. This work was partially supported by Deutsche Forschungsgemeinschaft (grant CRC 1310, to ML).

## Materials and methods

### Antigen-receptor interaction models

Here we model the binding (free) energy between an antigen with epitope sequence **a** = (*a*_1_, …, *a*_*ℓ*_) and a B-cell receptor (BCR) sequence **b** = (*b*_1_, …, *b*_*ℓ*_) as an additive function,

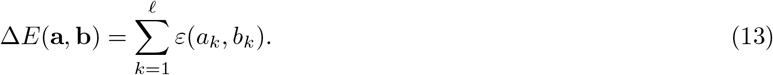

This function includes entropy contributions of non-translational degrees of freedom; i.e., rotations and elastic deformations of the molecules involved. We use three established models of amino acid interactions *ε*(*a, b*): the TCRec model originally inferred for T cell receptors [30], the Miyazawa-Jernigan matrix [31], and normally distributed random energies. In each case, we introduce a scale factor that is inferred from measured BCR-antigen binding energies (see below). The zero point of Δ*E*, by definition, corresponds to a reference antigen-BCR pair with equilibrium binding constant *K*_0_ = 1M. In this gauge, the binding energy and the dissociation constant *K* of an arbitrary pair are related by

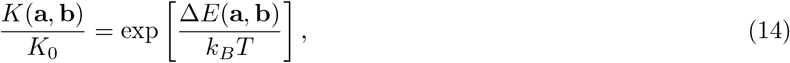

where *k*_*B*_ is Boltzmann’s constant. In the main text and below, we express all energies in units of *k*_*B*_*T* at a fixed, physiological temperature *T*.

### Density of BCR states

To characterise the naive B-cell repertoire available for response to a given antigen **a**, we use the density of lineage states defined by a unique BCR sequence **b**,

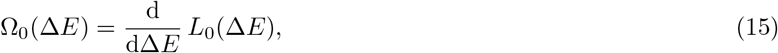

where *L*_0_(Δ*E*) is the expected number of lineages in an individual with binding constant *K* < exp(Δ*E*) to the epitope **a** (we suppress the dependence on **a** in this paragraph). By equation (14), this form is equivalent to the definition given in the main text, Ω_0_(*K*) = (*K*d/d*K*)*L*_0_(*K*). We further define the micro-canonical entropy

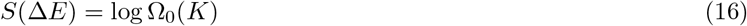

and the associated micro-canonical inverse temperature,

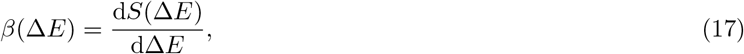

a parameter that is independent of the physiological temperature *T* appearing in equation (14). Because we measure energies in units of *k*_*B*_*T*, the parameter *β* gives the inverse temperature in units of (*k*_*B*_*T*)^−1^. To compute these micro-canonical quantities, we evaluate the canonical partition function,

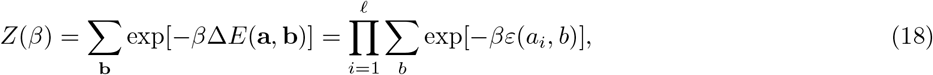

which depends on *β* as an independent parameter. This function defines the canonical binding energy,

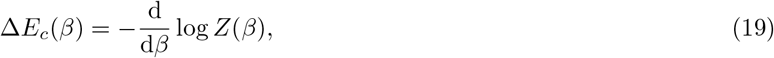

which is an expectation value in the ensemble (18), and the associated entropy,

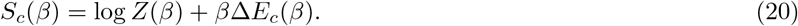

We invert the relation (19) to write the inverse temperature as a function of the binding energy, *β*(Δ*E*_*c*_), and we substitute this function into equation (20) to obtain *S*_*c*_(Δ*E*_*c*_). Upon equating Δ*E* = Δ*E*_*c*_, these functions provide an excellent approximation to their micro-canonical counterparts *β*(Δ*E*) and *S*(Δ*E*), as given by equations (16) and (17). Thus, the canonical formalism provides an efficient way to compute the density of states, Ω_0_(Δ*E*), for the system at hand.

### Antigen-receptor ensembles

To compare the response repertoires for different antigens, we evaluate the BCR lineage density Ω_0_(*K*, **a**) for a random sample of epitope sequences **a**. For a given amino acid interaction matrix and a given antigen, our energy model has two free parameters, the binding length *ℓ* and the scale factor of the energy, which sets the energy variance 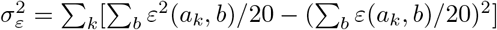. Here we calibrate these parameters by tuning the minimum binding constant expected in an individual repertoire and the global minimum binding constant to observed values of typical high-affinity antibodies generated in primary infections and of ultra-potent antibodies, *K*\* ≈ 10^−7^M and *K*_*m*_ ≈ 10^−11^M [32, 1]

The resulting ensemble of response repertoires has the following properties (Fig. S2): (i) The distributions of inferred binding lengths and of the rms. energy variation per site are strongly peaked around values *ℓ* ∼ 20 and *Σ*_*ε*_/*ℓ*^1/2^ ∼ 1. (ii) A higher energy variance per site can be traded for a shorter binding length, consistent with a constraint on the total energy variance 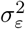. (iii) The lineage densities *ρ*_0_(*K*) depend only weakly on the antigen sequence **a** and have similar inverse activation temperatures, *β*\* ≡ *β*(*K*\*) = 2.5 *±* 0.3.

### Activation of B-cells by kinetic proofreading

An early infection is characterised by low densities of antigens and B-cells. Accordingly, we model the activation of individual B-cells upon functional binding with a single antigen. We assume that activation of antigen-bound cells requires a chain of *p* irreversible steps with characteristic rate *k*_step_ (Fig. 1). Hence, the B-cell activation rate takes the form

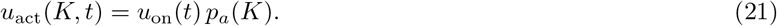

Here, the association rate per B-cell, *u*_on_(*t*) = *b*_0_*k*_on_*ρ*_*A*_(*t*) is proportional to the number of receptors per cell, *b*_0_, the diffusion-limited association rate to a single B-cell receptor, *k*_on_, and the antigen density, *ρ*_*A*_(*t*). The probability of activation after association is given by

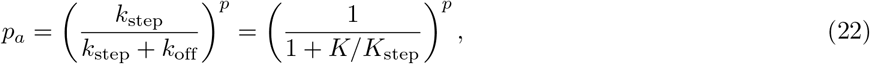

where *k*_off_ = *Kk*_on_ and *k*_step_ = *K*_step_*k*_on_. At each intermediate state, the antigen can dissociate or undergo the next activation step; these alternatives are independent Poisson processes with rates *k*_off_ and *k*_step_, respectively. Hence, the next activation step occurs before dissociation with probability *k*_step_/(*k*_step_ + *k*_off_).

The probability that a lineage gets activated up to time *t* is

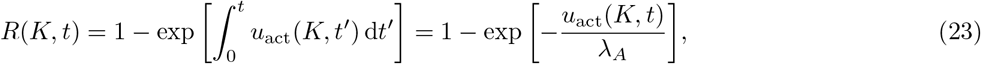

as given by equation (2) of the main text. Here we have used that antigens proliferate exponentially, *ρ*_*A*_(*t*) ∼ exp(*λ*_*A*_*t*). At early times, activation is association-limited and rare for all *K*,

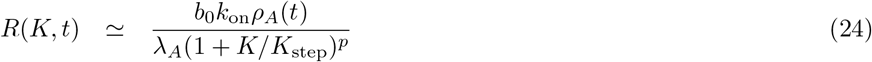

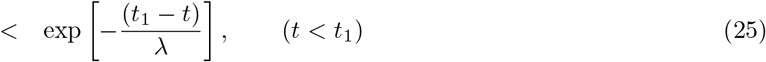

where

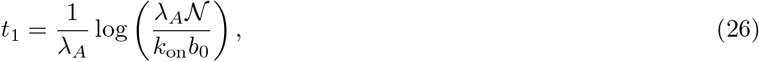

and 𝒩 is Avogadro’s number. For *t* > *t*_1_, we have

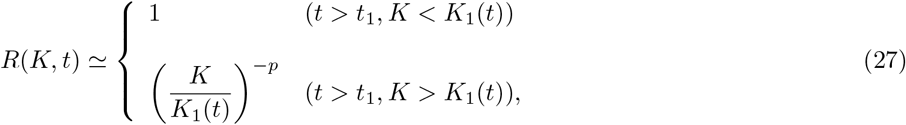

with

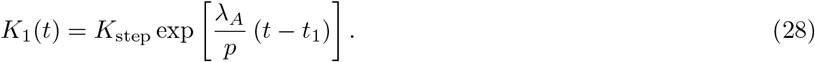

That is, deterministic activation of B-cell lineages occurs along a moving front, *K*_1_(*t*). Ahead of the front, activation is strongly suppressed by proofreading.

### Alternative models of B-cell activation

To highlight the specific role of kinetic proofreading in the activation of naive B-cells, we compare the proposed activation mechanism to alternative mechanisms with reversible binding kinetics (Fig. S1). The corresponding rates govern transitions between unbound and intermediate antigen-bound B-cell states (marked by grey shading in Fig. 1 and Fig. S1). We note that all activation mechanisms have at least one irreversible step: the last transition to exponential proliferation and antibody production (marked by green shading).

In an early primary infection, the activation of B-cells takes place under specific physiological conditions: (i) The antigen density and, hence, the equilibrium occupancy of B-cells remains low (*ρ*_*A*_/*K* ≲ 10^−1^, given *ρ*_*A*_ ≈10^11^/l ≲ 10^−13^M and *K* ≳ *K*\* ≈ 10^−7^M). These conditions differ drastically from the densities in confined spaces, e.g., lymph nodes and germinal centers, which are relevant for the binding kinetics of presented antigens. (ii) The association kinetics of virions and plasma B-cells is believed to be diffusion-limited; i.e., it takes place at a homogeneous rate *k*_on_ [46]. This excludes mechanisms for specific recognition by formation of immunological synapses, which have been proposed for presented antigen and operate by modulation of an activation-limited rate *k*_on_ [47, 48, 49, 17, 50]. (iii) For efficient activation, the rate *k*_step_ cannot be smaller than all other transition rates, as it is usually assumed in models of kinetic proofreading [14, 15]. Tuned rates discussed in the main text are of order *k*_step_ = 4 · 10^−3^s^−1^, which implies *k*_step_ ≳ *k*_off_ for high-affinity B-cell lineages. In this regime, the activation dynamics at a given antigen density *ρ*_*A*_ is close to a non-equilibrium steady state, even if the binding kinetics satisfies detailed balance. We consider two specific classes of models with diffusion-limited association and reversible binding kinetics:

- Activation via an excited intermediate state. This process is a reversible analogue of the proofreading dynamics discussed in the main text. Allowed transitions are between the unbound state and the primary bound state (with rates *k*_on_, *k*_off_) and between the primary bound state and the excited intermediate state (with rates 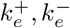), as shown in Fig. S1. Using detailed balance, the antigen-bound states have reduced binding energies Δ*E* = log(*K/K*_0_) and Δ*E*_*e*_ = Δ*E* + ΔΔ*E*_*e*_ respectively, where *K* = *k*_off_/*k*_on_ and 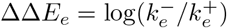). Like the proofreading model, this reversible model has a deterministic activation front given by equations (26) and (28). However, the activation probability is asymptotically proportional to the equilibrium occupancy of the intermediate state,

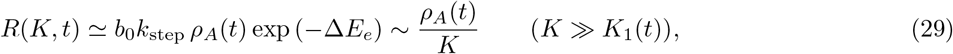

leading to weak suppression of low-affinity lineages [14, 15].
- Activation by cooperative binding. In this process, activation requires binding of two or more virions to receptors of the same B-cell, which has been observed for antigens actively transported to lymph nodes and presented to B-cells [48, 49, 51]. A bound state of *p* virions has the reduced binding energy Δ*E*_*p*_ = (*p*Δ*E* + ΔΔ*E*_*p*_) = log(*K*_*p*_/*K*_0_), where Δ*E*_*p*_ = log(*K/K*_0_) is the single-particle binding energy and ΔΔ*E*_*p*_ is the contribution of cooperative binding. Fig. S1 shows the case *p* = 2, where ΔΔ*E*_*p*_ = log(*k*′_off_/*k*_off_). In the cooperative binding model with detailed balance, the asymptotic activation probability is proportional to the equilibrium occupancy of the *p*-virion bound state,

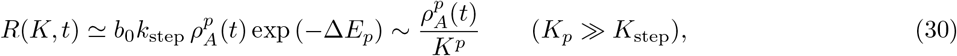

with *K*_step_ = *k*_step_/*k*_on_. For *p* > 1, this model leads to stronger suppression of low-affinity lineages; however, at the low antigen concentrations of an early infection, even high-affinity naive lineages do not reach deterministic activation (*R* ≪ 1 for *ρ*_*A*_(*t*)/*K* ≲ 10^−1^).

We conclude that the kinetic proofreading mechanism of B-cell activation introduced in the main text is the simplest model to generate deterministic activation of high-affinity lineages together with strong suppression of low-affinity lineages under the physiological conditions of an early primary infection.

### Activation statistics of B-cell repertoires

Given the density of naive B-cell lineages, Ω_0_(*K*), and the activation probability *R*(*t, K*), we can evaluate the density of activated lineages,

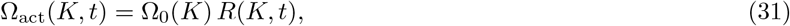

and the total number of activated lineages, *L*_act_(*t*) = ∫ Ω_act_(*K, t*) d(log *K*). Two activation regimes emerge:

- Low-specificity (LS) regime. In this regime, the specificity of activation is limited by the number of proofreading steps. According to equation (24), the function Ω_act_ is strongly peaked at a value *K*_*p*_ defined by the condition *β*(*K*_*p*_) = *p* (Fig. 3A). Integrating this function yields the expected number of activated lineages at time *t*,

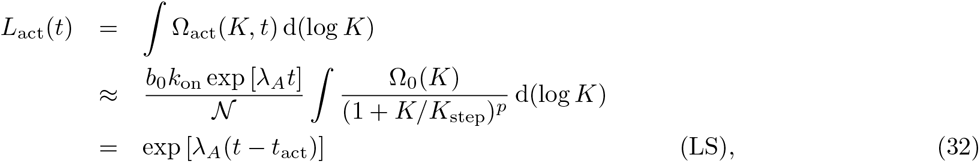

where *t*_act_ is given by the condition *L*_act_(*t*_act_) = 1.
- High-specificity (HS) regime. In this regime, the specificity of activation is limited by the complexity of the naive repertoire. According to equation (27), the function Ω_act_ is peaked around the moving front, *K*_1_(*t*) (Fig. 3B). In this case, we obtain

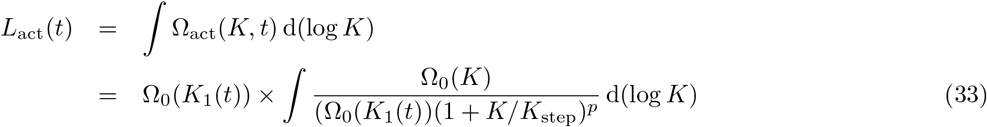

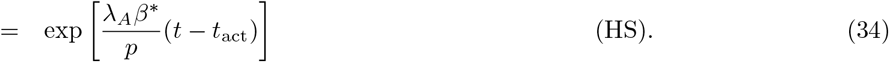

Here we use equations (16) and (28), and we note that the integrand in (33) has a peak value of order 1 and depends only weakly on *t*.

These results are given in equations (6) and (7) of the main text. In the activation dynamics discussed here, we assume that genetically distinct B cell lineages are also distinguishable in terms of their antigen binding affinity. Specifically, in our energy models, random mutations generate a binding energy change of order *k*_*B*_*T*. If the sequence-energy map is highly degenerate, multiple activations occurring in sequence clusters of very similar antigen affinity can generate new scaling regimes.

### Statistics of clone size

Here we compute the cumulative distribution function (CDF) of clone size, Φ(*N, t*), which is defined as the fraction of activated clones with size > *N* at time *t*. Given exponential growth with rate *λ*_*B*_, this function is given by the fraction of lineages activated before a time *t*′(*N*),

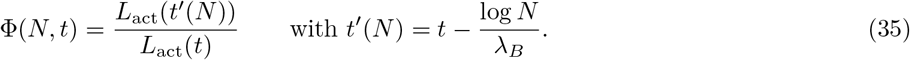

Using equations (32) and (34), we obtain

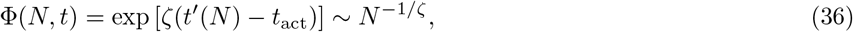

where the exponent *ζ* is defined in equation (10) of the main text. In a similar way, we compute the expected size of the *j*-th largest clone, ⟨*N*_*j*_⟩(*t*) (*j* = 1, 2, 3, …). We write ⟨*N*_*j*_⟩(*t*) ∼ exp[*λ*_*B*_(*t* − *t*_*j*_)], where the activation time *t*_*j*_ is given by the condition *L*_act_(*t*_*j*_) = *j*. Using again equations (32) and (34), we have *t*_*j*_ ∼ *ζ* log *j* and

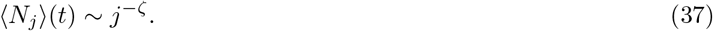

Equations (36) and (37) are related by Zipf’s law. Both are independent of *t* and, hence, valid also for the saturation clone sizes, 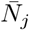, as used in equation (9) of the main text.

### Empirical clone size statistics in early germinal centers

We analyze sequencing data of early germinal centers (GCs) from Tas et al. [11] as a proxy for the initial population of activated B-cell clones. We use CDR3 sequence counts to estimate the clone size of the different B-cell lineages taking part in the response. The dataset contains samples from 6 dissected lymph nodes, each of which contains 2 GCs. We calculate the expected scaled clone size of the *j*-th largest clone found in each lymph node, *N*_*j*_/*N*_1_ (thin lines in Fig. 4A, insert) and the average value over all 6 lymph nodes (thick line in Fig. 4A, insert). We also calculate the clonal entropy Σ = − Σ_*j*_ *x*_*j*_ log *x*_*j*_ with *x*_*j*_ = *N*_*j*_/*N* and *N* = Σ_*j*_ *N*_*j*_ (the sum runs over all clones in the dataset).

### Statistics of antigen affinity

We now evaluate the CDF of antigen binding constants, Φ(*K, t*) = *L*_act_(*K, t*)/*L*_act_(*t*), in the HS regime. For the high-affinity tail of this function, *K* ≳ *K*\*, activation occurs deterministically, which implies *L*_act_(*K, t*) ≃ *L*_0_(*K*). Recalling that Ω_0_(*K*) = (*K*d/d*K*)*L*_0_(*K*), we obtain Φ(*K*) ∼ *L*_0_(*K*) ∼ Ω_0_(*K*). Hence,

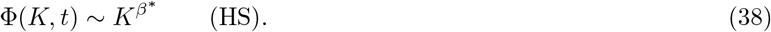

Using Zipf’s law, as for the clone size, we obtain the expectation value of the *l*-th lowest binding constant,

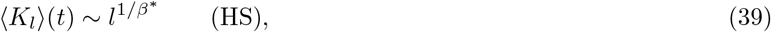

as given in equation (11) of the main text. Again, this relation is independent of *t* and valid also at the saturation point. Because activation occurs on the moving front *K*_1_(*t*), the clone rankings by size and affinity are equivalent up to fluctuations. Hence, by combining equations (37) and (39), we obtain a power law relating size and affinity

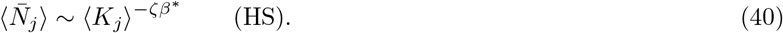

In the LS regime, there is no clear power law relation between affinity and rank (Fig. 5B). High-affinity activated clones span the range between *K*\* and *K*_*p*_ and show a faster decline of affinity with rank than in the HS regime (Fig. 5BC). Hence, empirical exponents fitted to affinity-rank data take values > 1/*β*\* (Fig. 5D).

### Potency statistics and elite neutralisers

In the HS regime, equations (37) and (39) also determine the statistics of single-clone potencies 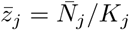,

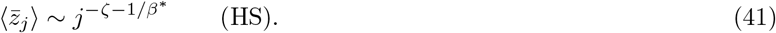

For typical individuals, many clones contribute to the total potency 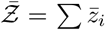. However, a characteristic of Luria-Delbrück-Delbrück models is the existence of giant fluctuations. In the HS regime of the model proposed here, there is a set of elite neutralisers singled out by early activation of their first clone. These “jackpot” clones have simultaneously high affinity and large size, which takes a sizeable fraction of the total activated repertoire, 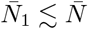 (Fig. 4D). The CDF of potency, Φ(*Ƶ*), is defined as the fraction of individuals with 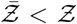. For 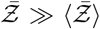, potency is dominated by jackpot clones, 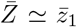. We find two scaling regimes (Fig. S4C). In the pre-asymptotic regime 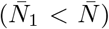, size fluctuations of the jackpot clone are dominant, and we can write 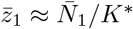. Hence, by equation (36), the CDF of potency takes the form

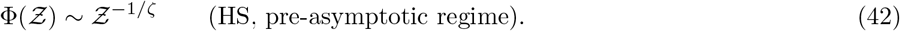

In the asymptotic regime 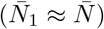, affinity fluctuations are dominant, and we have 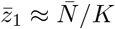. Hence,

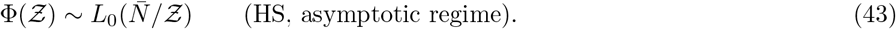

### Model parameters and numerical simulations

In the analytical and numerical analysis, we use the following empirical parameters for the activation process and the B-cell repertoire: (i) Size of the naive repertoire: *L*_0_ = 10^9^ lineages [2, 4, 3, 1]. (ii) Growth rate of the antigen population: *λ*_*A*_ = 6d^−1^ [5, 6, 7, 8]. (iii) Proliferation rate of activated B cells: *λ*_*B*_ = 2d^−1^ [28]. (iv) Activation step rate: *k*_step_ = 0.5min^−1^. As discussed in the main text, this value is tuned to a repertoire size *L*_0_ = 10^9^. (v) Antigen-BCR diffusion-limited association rate, *k*_on_ = 10^6^M^−1^*s*^−1^ [27]. (vi) Carrying capacity of activated B cells: 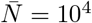 cells [28]. In the optimality analysis of tuned repertoires Fig. S3, we vary *p, k*_step_, and *L*_0_ around the LS-HS crossover point given by *p* ∼ *β*\*(*L*_0_), *k*_step_ ≈ *k*_on_*K*\*(*L*_0_).

To simulate a primary B-cell response, we start by creating an initial population of *L*_0_ B-cell lineages. Each B-cell lineage is represented by a BCR sequence of length *ℓ*, randomly drawn with the set of 20 amino acids. Assuming a deterministic expanding antigen concentration as defined in the main text, we calculate the non-homogeneous activation rate of each B-cell lineage, *u*_act_(*t, K*), as given in equation 21. Here we use the fact that the time for the first event, *t*_1_, in a non-homogeneous Poisson process with rate *u*(*t*) is distributed according to

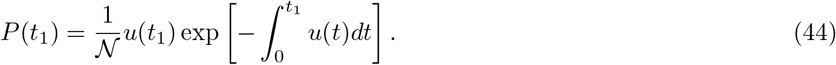

with 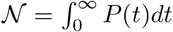. We sample then the activation time of each B-cell lineage by sampling uniformly distributed random numbers *r* ∼ 𝒰[0, 1] and then using the inverse of the cumulative version of equation 44. Once we have all activation times, we proceed to determine the clone size of each of the B-cell clones. Here we integrate *L*_act_ coupled differential equations

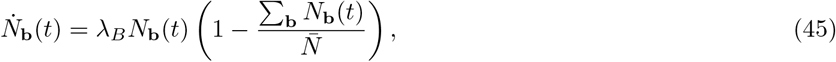

assuming all clones start with clone size *N*_**b**_(0) = 1. We neglect all B-cell clones whose final clone size is smaller than 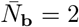, corresponding to less than one cell division. All notebooks, codes and input files used in this work are available at https://github.com/rmorantovar/primary_immune_response.

## Supplementary Figures

**Fig. S 1.**
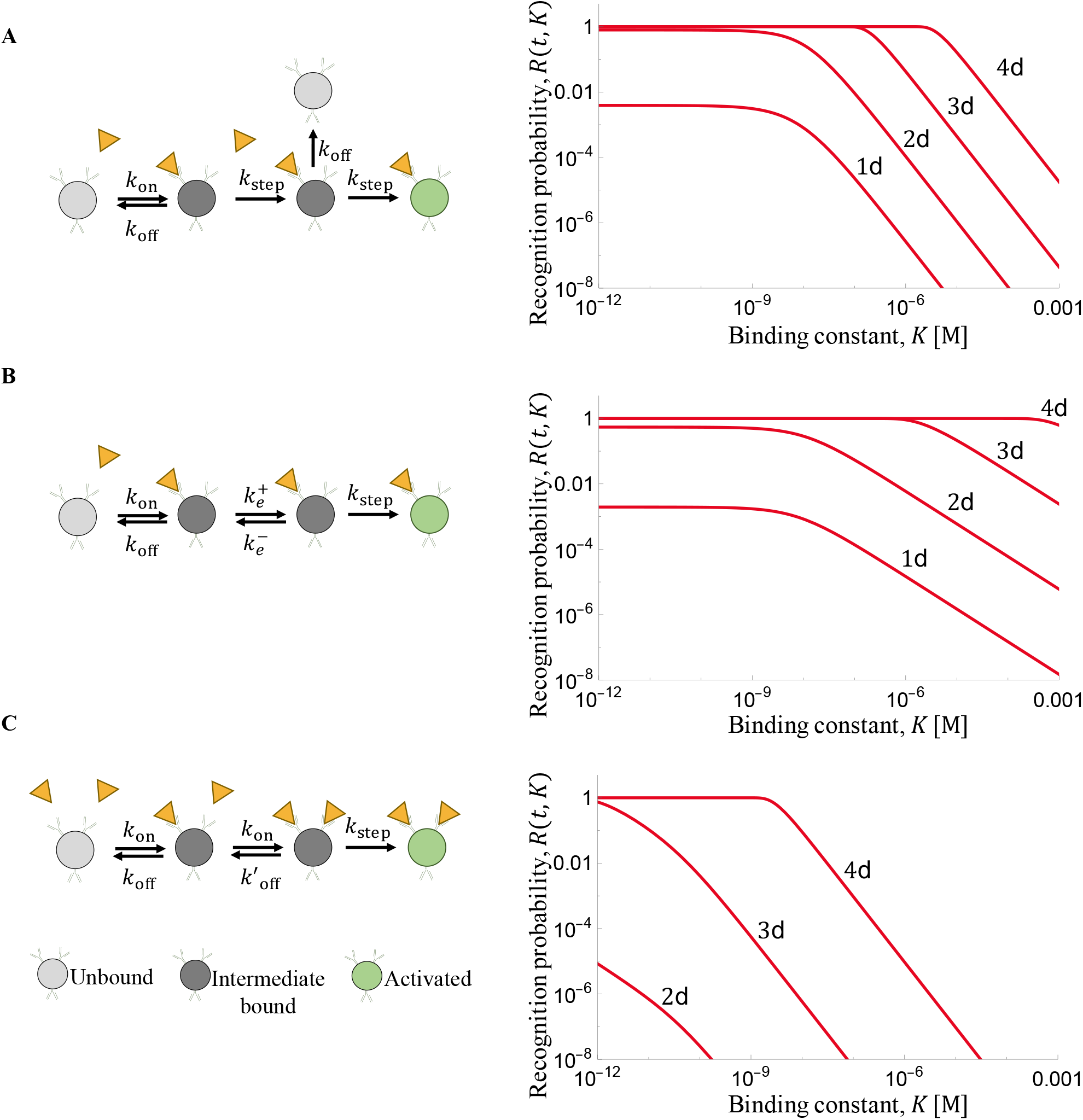
Mechanisms of B-cell activation. A schematic of the activation process (left) and the recognition probability *R*(*t, K*) (*t* = 1d, …, 4d after the start of the infection) are shown for the kinetic proofreading model discussed in the main text and for two models with reversible binding kinetics. **(A)** Activation by kinetic proofreading (as in Fig. 1). This process has *p* intermediate bound states and *p* irreversible activation steps (*p* > 1, shown here for *p* = 2). For antigen affinities of typical naive B-cells (*K* ≳10^−7^*M*), kinetic proofreading generates fast, deterministic recognition of high-affinity lineages (*R* ≃ 1) and strong suppression of low-affinity lineages (*R* ∼ *K*^−*p*^ for *K* ≫ *K*_1_(*t*)). **(B)** Activation via an intermediate excited state. This process is similar to the kinetic proofreading model, but the binding kinetics is reversible. It generates fast, deterministic of high-affinity lineages but only weak suppression of low-affinity lineages (*R* ∼ *K*^−1^ for *K* ≫ *K*_1_(*t*)). **(C)** Activation by cooperative binding. In this process, activation requires simultaneous binding of *p* virions to the same B-cell (*p* > 1, shown here for *p* = 2). It generates strong suppression of low-affinity lineages (*R* ∼ *K*^−*p*^ for *K* ≫ *K*_step_), but even high-affinity naive lineages do not reach deterministic activation (*R* ≪ 1 for *K* ∼ 10^−7^*M*). Kinetic parameters: *k*_on_ = 10^6^*s*^−1^*M* ^−1^; *’
s* = 10^−2^ ; *k*_step_ = 0.5min^−1^; 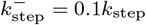.

**Fig. S 2.**
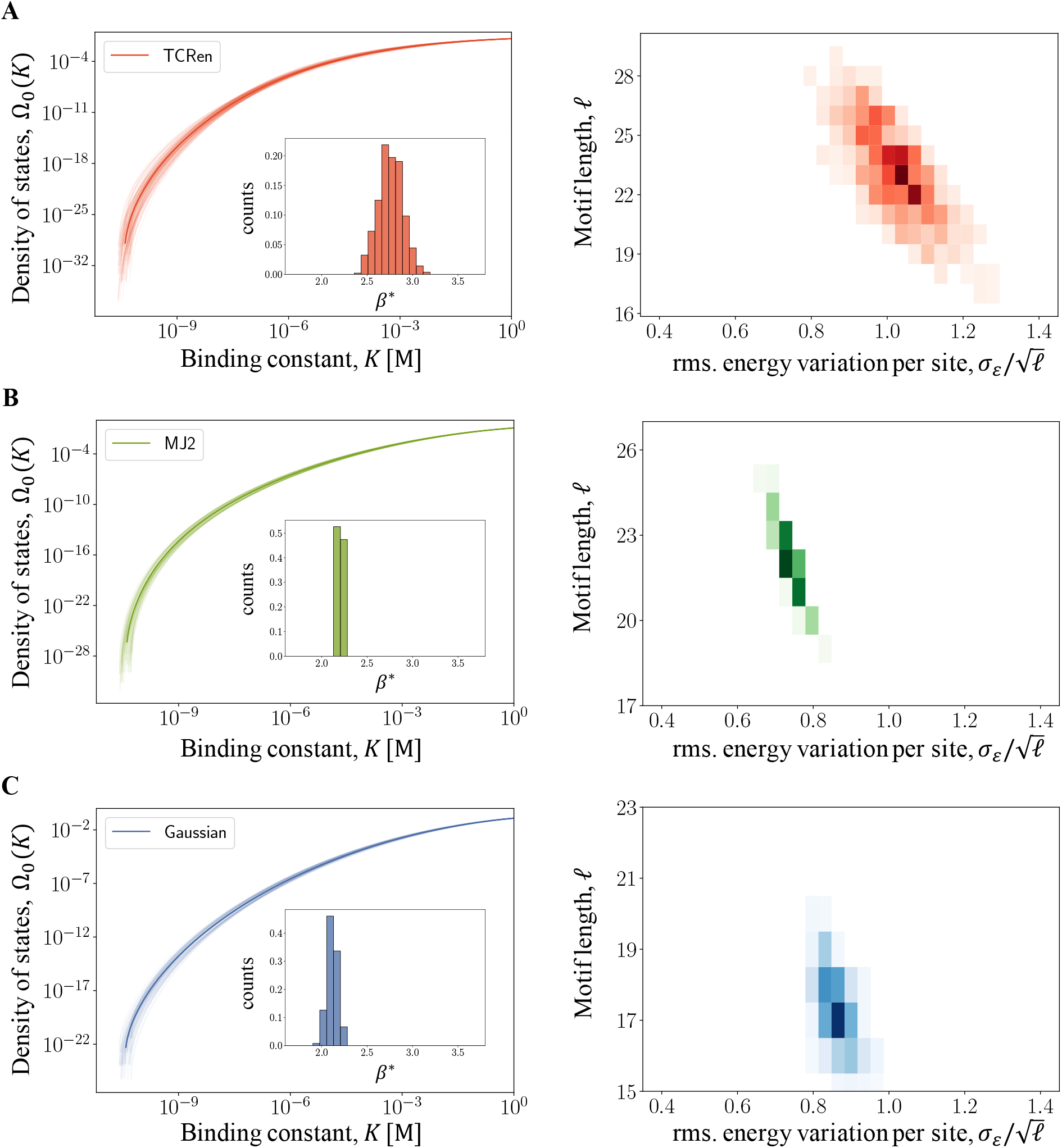
Density of states in different energy models. Left: density of naive B-cell lineages, Ω_0_(*K*), for a sample of antigen epitope sequences with *K*_*m*_ ≈ 10^−11^*M* and *K*\* ≈ 10^−7^*M* (thin lines: individual antigens, thick line: ensemble average); see SI text. Insert: ensemble distribution of the inverse activation temperature *β*\*, as defined in the main text. Right: joint distribution of the binding length, *ℓ*, and of the rms. energy variation per site, *σ*_*ε*_/*ℓ*^1/2^. Binding motifs are randomly sampled from different additive energy models: **(A)** TCRen matrix [30], **(B)** Miyazawa-Jernigan matrix [31], **(C)** Gaussian random energies; details are given in SI Text. Model parameters as in Fig. 2.

**Fig. S 3.**
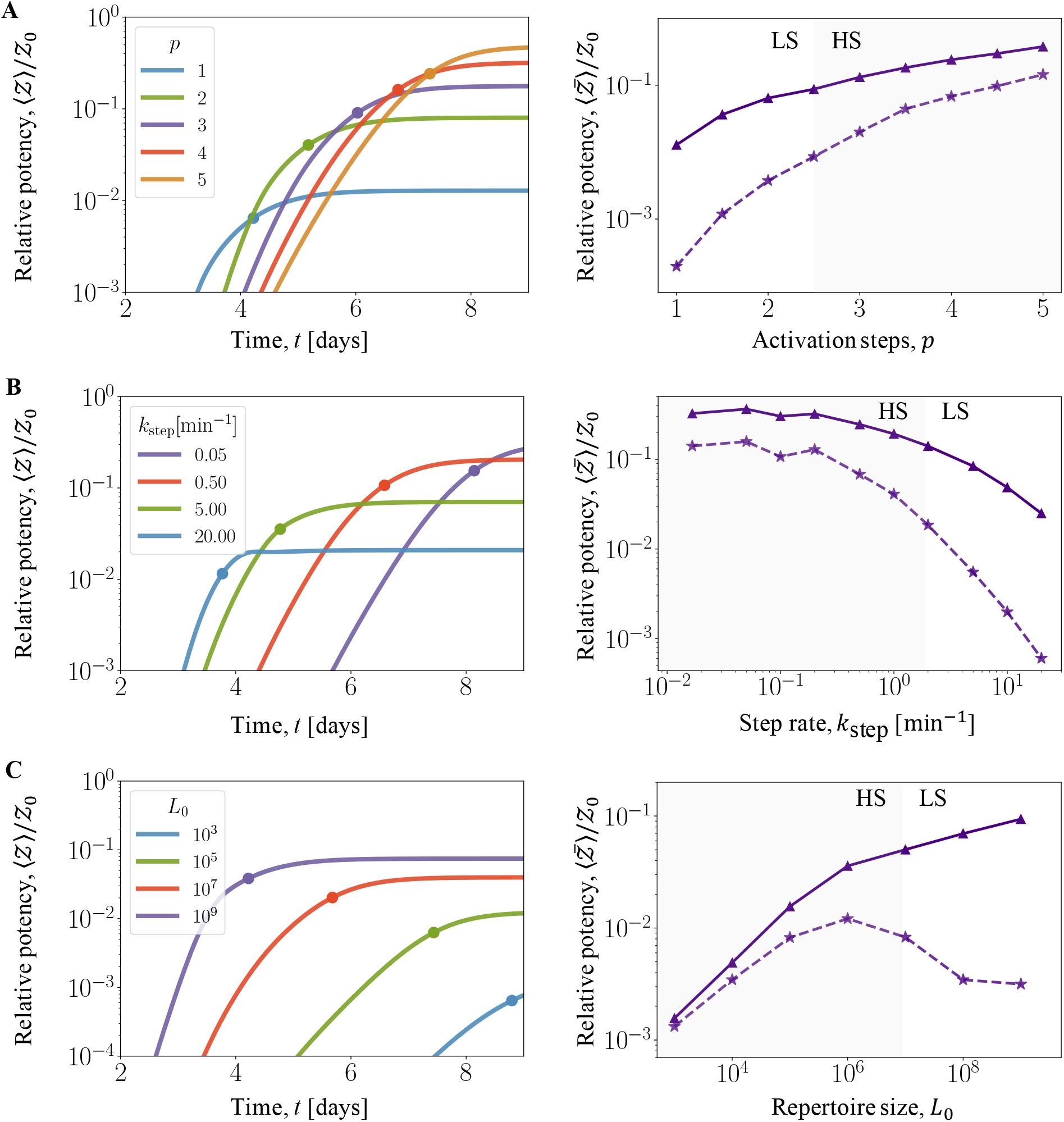
Optimality of antigen recognition by tuned proofreading. Antiserum potency is shown for variation of the proofreading parameters *p, k*_step_, and of the repertoire size, *L*_0_, around the LS-HS crossover point (*p* ∼ *β*\*(*L*_0_), *k*_step_ ≈ *k*_on_*K*\*(*L*_0_), *L*_0_ = 10^9^), which characterises a tuned repertoire as discussed in the main text. Left: time-dependent, population averaged potency, *Ƶ*(*t*); the half-saturation point (*t*_50_, *Z*_50_) is marked by a dot. Right: saturation value, 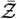 (thick lines), and the potency component of the largest clone, 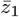 (dashed lines). All potencies are shown in units of the reference value *Ƶ*_0_ defined in the main text; the HS regime is marked by shading. **(A)** Variation of the number of proofreading steps, *p*, at constant *k*_step_ and *L*_0_ (cf. Fig. 4EF). In the LS regime (*p* ≲ *β*\*), potency increases strongly with *p*. In the HS regime (*p* ≳ *β*\*), increasing *p* yields a diminishing return of potency, while *t*_50_ continues to increase proportionally to *p*. **(B)** Variation of the rate of proofreading steps, *k*_step_, at constant *p* and *L*_0_. In the LS regime (*k*_step_ ≳ *k*_on_*K*\*), potency increases strongly with decreasing *k*_step_. In the HS regime (*k*_step_ ≲*k*_on_*K*\*), decreasing *k*_step_ yields a diminishing return of potency, while *t*_50_ continues to increase proportionally to 1/*k*_step_. **(C)** Variation of the repertoire size, *L*_0_, at constant *p* and *k*_step_. Here we use *p* = 2 and *k*_step_ = 0.05min^−1^, which are tuned for *L*_0_ = 10^7^. In the HS regime (*L*_0_ ≲ 10^7^), potency increases strongly with *L*_0_. In the LS regime (*L*_0_ ≳ 10^7^), increasing *L*_0_ yields a diminishing return of potency and the potency component of the largest clone decreases. Other model parameters as in Fig. 2.

**Fig. S 4.**
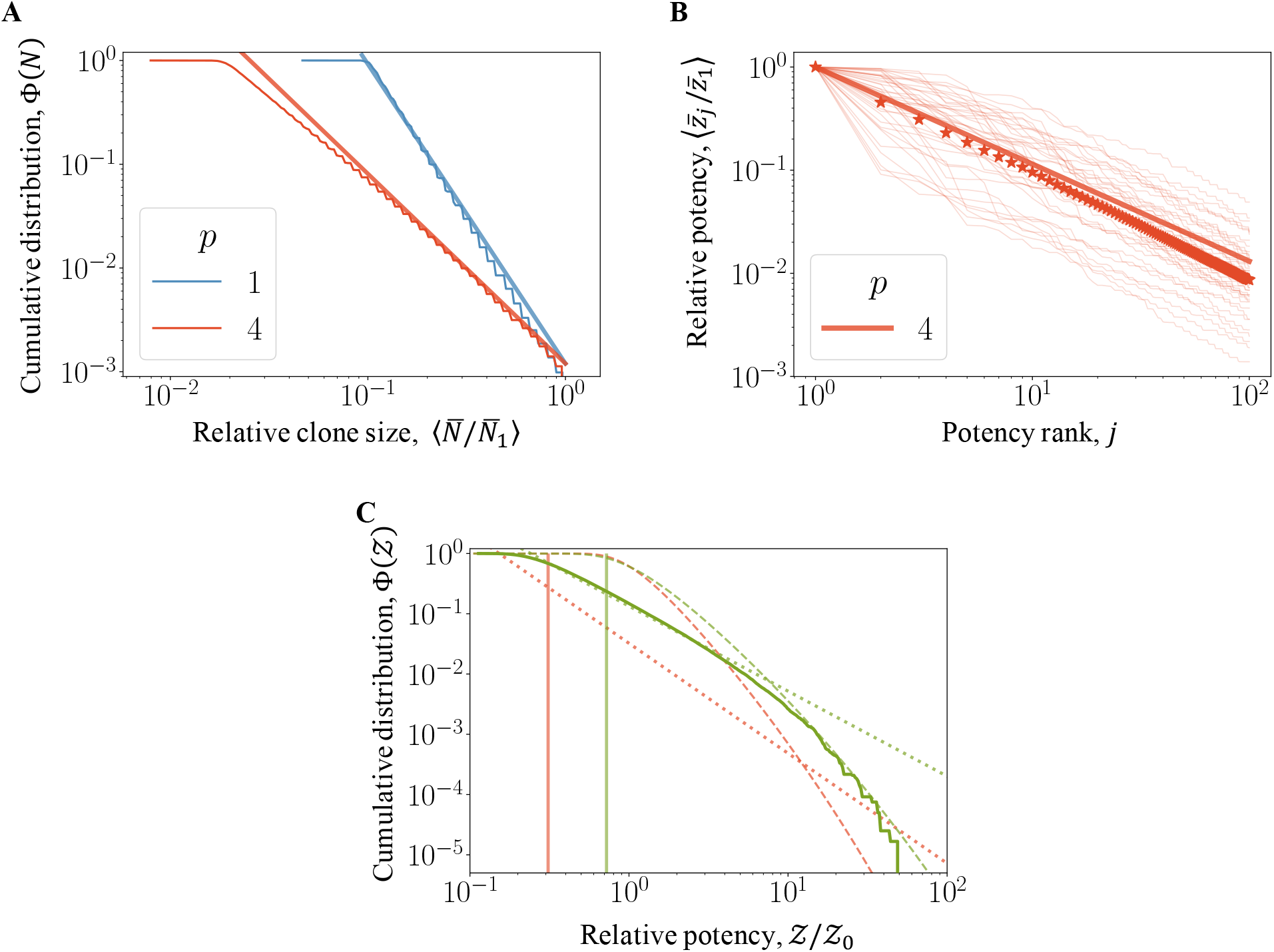
Statistics of activated repertoires. **(A)** Cumulative clone size distribution aggregated over individuals, Φ(*N*) ∼ *N* ^−1/*ζ*^ (blue: LS regime, *p* = 1; red: HS regime, *p* = 3). **(B)** Potency-rank relation in the HS regime, 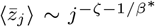 for potency-ranked clones (*j* = 1, …, 100). Thin lines give rankings in a set of randomly chosen individuals. **(C)** Cumulative population distribution function of potency in the HS regime, Φ(*Ƶ*) (*p* = 4; *L*_0_ = 10^7^ in green and *L*_0_ = 10^9^ in red). Elite neutralisers with jackpot clones define a pre-asymptotic regime, Φ(*Ƶ*) ∼ *Ƶ*^−1/*ζ*^ (dotted), and an asymptotic regime, 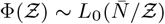 (dashed). Vertical lines indicate population average 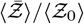. Model parameters as in Fig. 2.

